# Motor experience leads to structural changes in the adult male mouse inferior olive

**DOI:** 10.64898/2026.09.03.747181

**Authors:** Tara H. Turkki, Sophia Dubois, Hugo Hoedemaker, Tetiana Salamovska, Bogna M. Ignatowska-Jankowska, Marylka Y. Uusisaari

**Affiliations:** Neuronal Rhythms in Movement Unit, Okinawa Institute of Science and Technology, 1919-1 Tancha, Onna-son, 904-0495, Okinawa, Japan; Neural Computation Unit, Okinawa Institute of Science and Technology, 1919-1 Tancha, Onna-son, 904-0495, Okinawa, Japan; Computational Neuroscience Unit, Okinawa Institute of Science and Technology, 1919-1 Tancha, Onna-son, 904-0495, Okinawa, Japan

**Keywords:** inferior olive, dendritic morphology, gap junctions, cerebellum, motor learning

## Abstract

The activity of the inferior olive (IO), conveyed to the cerebellar cortex via climbing fibers and manifesting as the complex spikes (CSs) in Purkinje neurons, is at the core of major theories of cerebellar function and motor learning. While the computational meaning of the CSs remains under intense debate, it is well accepted that changes in their occurrence and timing play a critical role in guiding cerebellar plasticity. However, it has not been examined whether the plastic processes leading to these changes in cerebellar CS activity are expressed in the IO, or whether they are entirely caused by changes in brain structures upstream of the climbing fibers, including the cerebellar nuclei.

Here, we examine whether prolonged, kinematically challenging motor experience can lead to detectable changes in the IO network structure that could underlie shifts in cerebellar CS activity. As the IO neurons communicate exclusively via electrical synapses (gap junctions) residing on dendritic structures, we hypothesized that if long-term increases in electrical coupling related to the insertion of new gap junctions occur in response to novel motor experiences, morphological changes in the dendrites would also be expected.

To investigate this, we quantitatively characterized the geometry of the IO neuropil revealed by immunohistochemical staining in adult male mice. We found that the neuropil structure varies across olivary subnuclei in naive animals, possibly underlying known differences in cerebellar complex spike co-activation patterns. Geometrical measures related to structural complexity also revealed that exposing the animals to a challenging high-speed treadmill running task requiring full-body coordination led to localized changes suggestive of increased network connectivity. In addition, the density of immunofluorescence puncta labeling Cx36 increased in the same regions, supporting the notion that exposure to contexts where novel motor skills need to be acquired may lead to changes in the clustering strength among IO neurons and thereby restructuring of the olivo-cerebellar micromodules. To our knowledge, this is the first report of experience-related plasticity within the IO, calling for renewed attention to the role of the IO in olivocerebellar function.

## Introduction

Current research devoted to understanding motor learning and cerebellar plasticity examines processes downstream of learning-induced changes in the timing of complex spikes (CSs) in the Purkinje neurons (PNs), but virtually nothing is known about plasticity at their source, the inferior olive (IO). As the IO neurons are proposed to “gate” the translation of afferent signals into these learning-guiding CSs^1–3^, it is necessary to examine whether the IO and its intrinsic processes also undergo changes during learning.

The IO lacks local chemical synapses, and instead is spatially organized into electrically-coupled clusters that align within the olivo-cerebellar microzones^4^. This organization is supported by the uniquely complex dendritic and spine characteristics^5–10^. The spine glomerulus, a structure where numerous GJ-coupled spines as well as GABAergic and glutamatergic axon terminals wrapped in a glial sheet form a compact point of communication with modifiable strength^10–12^. Indeed, the short-term modulation of electrical coupling strength by activation of glomerular axon terminals has been targeted by several experimental and computational studies^13–17^. The central role in these frameworks is played by the GABAergic nucleo-olivary (NO) feedback pathway that originates in the contralateral cerebellar nuclei^18–20^. By its GABAergic nature, the NO pathway is proposed to allow the cerebellum to modulate its own complex spike firing, either by suppressing or desynchronizing olivary activity. While such effects can be demonstrated under experimental conditions^13,21^, the modulatory effects are rapidly reversed and hence unlikely to be the sole mechanism underlying long-term plasticity.

The IO receives numerous afferent, excitatory inputs such as head movement-related information through the superior colliculus^22,23^, oculomotor information through the medial accessory nucleus of Bechterew, the medial accessory oculomotor nucleus, the nucleus prepositus hypoglossi, and the nucleus of the optic tract^24,25^, balance from vestibular nuclei^26^, and cerebral information relay via the mesodiencephalic junction^27–30^. Learning or improving tasks dependent on cerebellar computation possibly involves axonal sprouting from relevant upstream sources in addition to conventional chemical synaptic plasticity, analogous to what has been observed in the cerebellar nuclei^31^. However, since single IO neurons are known to be inherently multimodal already in naive animals (e.g. responding both to auditory and tactile stimulations^32^ and capable of rapidly shifting their preferred modality in a context-dependent manner^33^), it seems unlikely that novel skill learning-related changes in complex spike activity could be entirely dependent on changes in afferent inputs.

What kind of intrinsic processes could underlie long-term plasticity of the IO neurons? The low, 1-2Hz firing rate of IO neurons has not been reported to change in any motor learning paradigms, suggesting that changes in expression of ion channels modulating intrinsic excitability are not a major factor.

Timing of individual IO spikes is controlled by the gap junction-dependent subthreshold oscillations^34–36^ and their co-activation may be higher among neurons with strong electrical coupling^37,38^. Hence, mechanisms that would allow long-term changes in the strength of coupling among IO neurons should be sought. As the small size of dendritic spines limits the potential for effective coupling strength increases by simple addition of connexin channels into the plaques, we hypothesized that significant strengthening of electrical coupling should be supported by the addition of neuropil structures to carry the added gap junctions. Such a need for a structural plasticity mechanism would align with the known slow timescale of motor learning (from days to weeks) as well as “permanence” of motor skill acquisition.

As the first step towards elucidating the mechanisms that could allow reshaping of the olivary clusters, we first describe the neuropil morphology in terms of overall density and segment-level curvature in naive mice based on skeletonizing confocal images where IO dendrites were labeled with MAP2-immunohistochemistry. The results demonstrate that the geometry of the IO neuropil is not homogeneous across the olivary subnuclei. Next, we assessed the effects of a 14-day motor training paradigm on the morphological complexity of the adult male mouse IO neuropil. In animals subjected to intense treadmill running, significant increases in neuropil complexity as well as gap-junctional labeling in the intermediate medial accessory olive (MAO), a region involved in balance-related motor functions.

## Results

### Two-week treadmill running experience paradigm

Across the 14-day treadmill training period (Figure 1A), the mice in the running group encouraged to run at their personal maximum speeds, the treadmill speed of the walking group mice was restricted to a maximum of 15 m/min, and the sedentary mice were not trained on the treadmill (Figure 1B,C). The exposure to running task was kept deliberately short to avoid overall body composition and cardiovascular fitness changes, and indeed the body weight or running mice only slightly trended behind the walking and sedentary groups(Figure 1D). All animals started from a similar initial speeds (14.75 *±* 0.87 and 14.44 *±* 1.62 m/min, for walker and runner animals, respectively). Throughout the training, walkers were not challenged with speeds above 15 m/min, while the runners were daily encouraged to run at increasing speeds. (Figure 1)E). By the end of the training, the mean speed reached by the runners was 34.50 *±* 5.70 m/min (Figure 1E). Running performance of individual mice is presented in Figure 1F as the fraction of trials where the mouse ran at *≥* 30 m/min on a given day, demonstrating diversity in performance, motivation, or capability across the training period.

**Figure 1.**
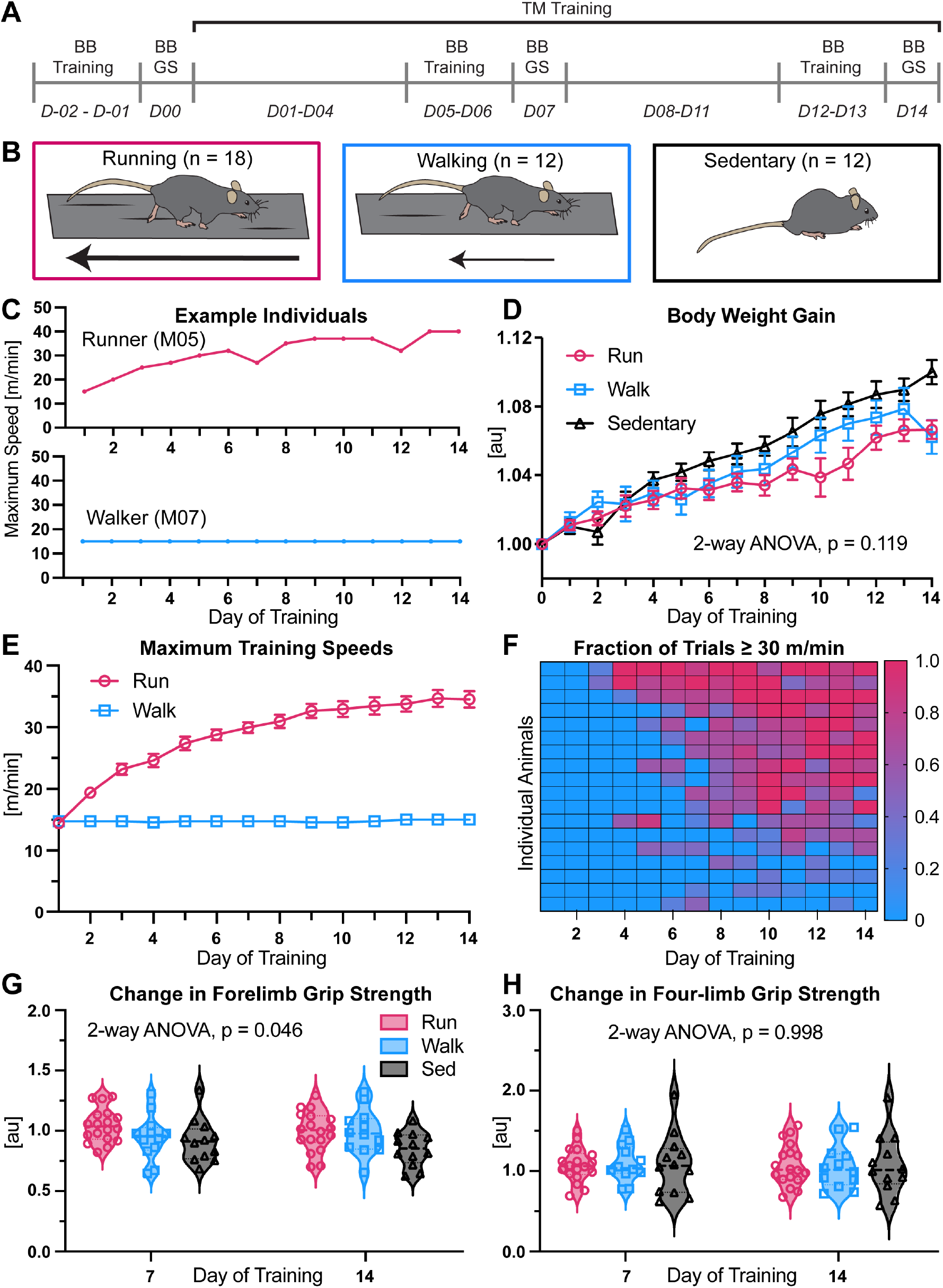
Motor experience in the form of high-speed treadmill running. (A) Experimental timeline for mouse behavior. (B) A schematic depiction of the three different groups for treadmill behavior in following experiments: running (30 m/min, left), walking (15 m/min, middle), sedentary (no treadmill behavior, right). (C) Examples of individual animal maximum speed trajectories across training. (D) Body weight measured on days 0-14 normalized to day 0. Two-way ANOVA with Geisser-Greenhouse correction for unequal variances. Data presented as mean *±* SEM. (E) Treadmill speeds used for the 14-day training. Data presented as mean *±* SEM. (F) Trials in which each individual mouse in the running group ran on a treadmill at a speed of 30 m/min or above out of the total trials on each training day. (G-H) Body weight and baseline-normalized forelimb and four-limb grip strength measured on days 7 and 14 of treadmill training.

A significant effect of group on forelimb grip strength was found (two-way ANOVA, F(2, 39) = 3.34, p = 0.046, Figure 1G). For the running group, forelimb grip strength increased from day 0 to day 7 (1.05 *±* 0.13 of baseline), while it decreased on day 14 (0.996 *±* 0.16 of baseline, Table 1). Meanwhile, in the walking and sedentary groups, grip strength on days 7 and 14 was lower than on day 0. While in the walking group, grip strength was 0.95 *±* 0.19 and 0.99 *±* 0.19 of baseline on days 7 and 14, respectively; the grip strength decreased more drastically in the sedentary group. Specifically on day 14, the sedentary group’s grip strength was only 0.85 *±* 0.14 of baseline (Figure 1G, Table 1). The post-hoc test revealed that the difference between the walking and sedentary group was significantly different in the change from baseline to day 7 (Table 1).

**Table 1.**
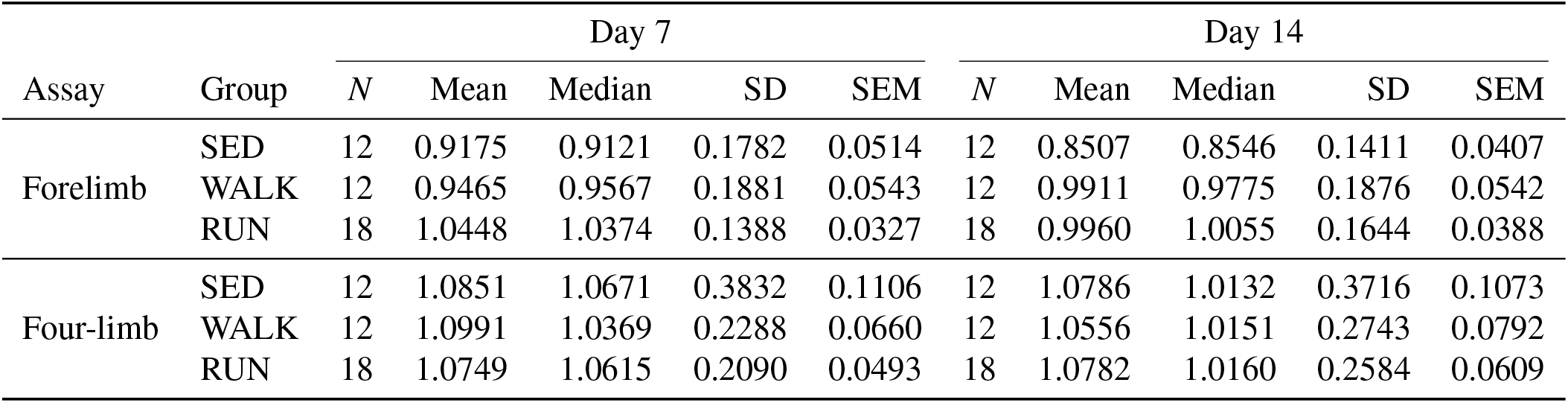
Mean, median, standard deviation (SD), and standard error of the mean (SEM) for body weight-normalized forelimb and four-limb grip strength relative to baseline (day 0) on days 7 and 14 of treadmill training.

| Assay | Group | Day 7 |  |  |  |  | Day 14 |  |  |  |  |
| --- | --- | --- | --- | --- | --- | --- | --- | --- | --- | --- | --- |
|  |  | *N* | Mean | Median | SD | SEM | *N* | Mean | Median | SD | SEM |
| Forelimb | SED | 12 | 0.9175 | 0.9121 | 0.1782 | 0.0514 | 12 | 0.8507 | 0.8546 | 0.1411 | 0.0407 |
|  | WALK | 12 | 0.9465 | 0.9567 | 0.1881 | 0.0543 | 12 | 0.9911 | 0.9775 | 0.1876 | 0.0542 |
|  | RUN | 18 | 1.0448 | 1.0374 | 0.1388 | 0.0327 | 18 | 0.9960 | 1.0055 | 0.1644 | 0.0388 |
| Four-limb | SED | 12 | 1.0851 | 1.0671 | 0.3832 | 0.1106 | 12 | 1.0786 | 1.0132 | 0.3716 | 0.1073 |
|  | WALK | 12 | 1.0991 | 1.0369 | 0.2288 | 0.0660 | 12 | 1.0556 | 1.0151 | 0.2743 | 0.0792 |
|  | RUN | 18 | 1.0749 | 1.0615 | 0.2090 | 0.0493 | 18 | 1.0782 | 1.0160 | 0.2584 | 0.0609 |

In all groups, an increase in four-limb grip strength was observed on both days 7 and 14. The changes from day 0 to day 7 were 1.075 *±* 0.21, 1.099 *±* 0.23, and 1.085 *±* 0.38, for the running, walking, and sedentary groups, respectively (Figure 1H, Table 1). On day 14, the four-limb grip strength of the running group increased slightly to 1.078 *±* 0.26, while in the walking and sedentary groups, the change in grip strength was less than on day 7, with 1.056 *±* 0.27 and 1.079 *±* 0.37, respectively. The group had no significant effect on four-limb grip strength (one-way ANOVA, F(2, 39) = 0.002, p = 0.998, Figure 1H, Table 1).

### Limited effects of the treadmill training paradigm on balance beam crossing performance

On days 0, 7 and 14, the animals in all three groups were challenged with a 70-cm balance beam crossing task (Figure 2A, B) to examine whether their increased treadmill performance generalized to overall agility. Notably, the task is relatively easy for the animals, leaving little room for improvement. Traverse duration (crossing time) of the balance beam of the mice in the running group was almost identical to baseline on day 7 (0.998 *±* 0.27), but the mice crossed faster on day 14 (0.697 *±* 0.21, Figure 2C1, Table 2) in comparison to baseline. The walking group followed a similar trend, while in the sedentary group, a similar decrease was observed at both time points (0.866 *±* 0.34 on day 7, 0.845 *±* 0.33 on day 14, Figure 2C1, Table 2). These changes were not significant. The mean postural height measured using the centroid of the animals as they crossed the beam was lower for the running and walking groups on day 7 compared to day 0, while it was higher for the sedentary group (Figure 2C2). Nearing the significance threshold (one-way ANOVA, F = 2.93, p = 0.065), this trend did not persist on day 14 (Figure 2C2). For mice in the running group, the number of slips slightly increased from the baseline to day 7 (0.12 *±* 2.67), and they decreased on day 14 (-1.82 *±* 3.07, Figure 2C3, Table 2). While the number of slips also decreased in the walking and sedentary groups both on days 7 and 14, the effect of group on the number of slips was not significant (one-way ANOVA, F = 0.42, p = 0.66 for day 7, F = 1.15, p = 0.33 for day 14).

**Figure 2.**
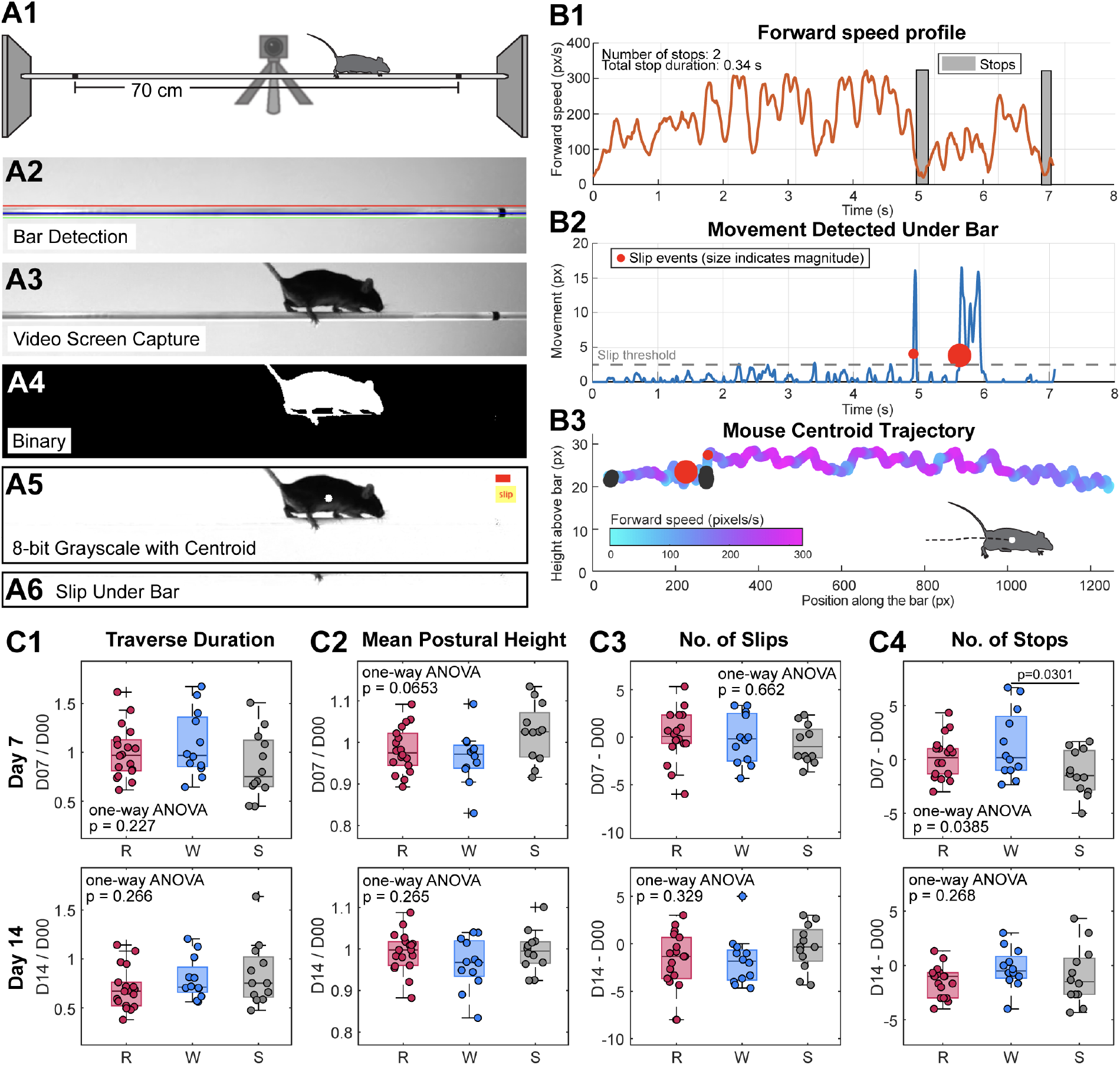
Motor experience has limited effects on balance beam crossing metrics. (A1) Schematic depiction of the balance beam setup, (A2–A6) workflow for video analysis to automatically detect beam crossing metrics. Example quantifications of (B1) forward speed and stop duration, (B2) detected movement under the bar and slip count, (B3) mouse centroid trajectory and instantaneous speed for a single trial. Change in (C1) traverse duration, (C2) mean postural height, (C3) the number of slips, and (C4) the number of stops from baseline (day 0). One-way ANOVA, Tukey’s HSD post-hoc test. *N* = 12–18/group.

**Table 2.** Mean, median, standard deviation (SD), and standard error of the mean (SEM) for the change in metrics of balance beam performance from day 0 to days 7 and 14 of treadmill training. Traverse duration and mean postural height were normalized to day 0 by division (ratio), whereas the numbers of slips and stops were normalized by subtraction (difference). *N* refers to the number of animals.

| Metric | Group | Day 7 |  |  |  |  | Day 14 |  |  |  |  |
| --- | --- | --- | --- | --- | --- | --- | --- | --- | --- | --- | --- |
|  |  | *N* | Mean | Median | SD | SEM | *N* | Mean | Median | SD | SEM |
| Traverse duration | SED | 12 | 0.866 | 0.752 | 0.336 | 0.097 | 12 | 0.845 | 0.752 | 0.325 | 0.094 |
|  | WALK | 12 | 1.084 | 0.969 | 0.336 | 0.097 | 12 | 0.795 | 0.711 | 0.213 | 0.061 |
|  | RUN | 18 | 0.998 | 0.970 | 0.266 | 0.063 | 18 | 0.697 | 0.671 | 0.214 | 0.050 |
| Mean postural height | SED | 12 | 1.023 | 1.026 | 0.070 | 0.020 | 12 | 0.995 | 0.995 | 0.049 | 0.014 |
|  | WALK | 12 | 0.964 | 0.972 | 0.063 | 0.018 | 12 | 0.964 | 0.968 | 0.062 | 0.018 |
|  | RUN | 18 | 0.982 | 0.975 | 0.055 | 0.013 | 18 | 0.993 | 0.998 | 0.048 | 0.011 |
| Number of slips | SED | 12 | −0.722 | −1.000 | 1.979 | 0.571 | 12 | −0.361 | −0.333 | 2.380 | 0.687 |
|  | Walk | 12 | −0.167 | −0.167 | 2.627 | 0.758 | 12 | −1.750 | −1.833 | 2.667 | 0.770 |
|  | RUN | 18 | 0.120 | 0.083 | 2.665 | 0.628 | 18 | −1.824 | −1.333 | 3.069 | 0.723 |
| Number of stops | SED | 12 | −1.250 | −1.500 | 2.085 | 0.602 | 12 | −0.944 | −1.500 | 2.685 | 0.775 |
|  | WALK | 12 | 1.333 | 0.167 | 3.175 | 0.917 | 12 | −0.250 | −0.500 | 1.826 | 0.527 |
|  | RUN | 18 | 0.194 | 0.167 | 1.924 | 0.453 | 18 | −1.454 | −1.000 | 1.405 | 0.331 |

The number of stops on day 7 increased in the running and walking groups from day 0, while they decreased in the sedentary group, and the effect of group was significant (one-way ANOVA, F = 3.55, p = 0.04, Figure 2C4). Specifically, a post-hoc pairwise comparison revealed that the difference between the walking and sedentary groups was significant (Tukey’s HSD, p = 0.03). On day 14, mice in all groups had a lower stop count than on day 0; however, this effect was not significant (one-way ANOVA, F = 1.36, p = 0.27, Figure 2C4).

### High-throughput imaging of the IO Neuropil

For high-throughput acquisition of MAP2-stained IO neuropil, automated generation of locations for high-resolution image stack (also referred to as region of interest, or ROI) acquisition was based on the estimated shape of the IO in the coronal section (Figure 2A1-A2). For each of the 9 slices containing the IO from each animal in the three groups, 25 of such image stacks were acquired(Figure 2A3-A4). Each image stack was examined and stacks that were out of focus or outside the IO were excluded before inclusion in the analysis. After the quality control step, 1037, 1239, and 1545 ROIs were included for the sedentary, walking, and running groups, respectively (Figure 2D).

The relative proportions of the subnuclei were similar across the three groups (15.82-17.27 % DAO, 35.59-38.06 % PO, and 45.76-47.14 % MAO, Figure 2D). Furthermore, once subdivided into anterior, intermediate, and posterior, the relative proportions also remained consistent across groups (Figure 2D). Very few ROIs were classified as posterior DAO and PO, and most of the posterior ROIs were represented by the MAO (89.46 % of all ROIs classified as “posterior” across subnuclei). Meanwhile, anterior ROIs made up approximately 40.24-42.70 %, 55.56-67.18 %, and 18.54-22.07 % of total DAO, PO, and MAO ROIs across groups, respectively, and the corresponding fractions of intermediate ROIs were 47.20-52.44 %, 30.78-39.91 %, and 28.08-29.79 % (Figure 2D). This demonstrates that the applied method produced high-throughput datasets with consistently classified ROIs across behavioral groups.

### Differences in the complexity of the IO neuropil in the naive mouse

Before investigating the possible effects of the treadmill training on neuropil structure, we examined the differences between IO subnuclei in the sedentary group. The individual CDFs representing the data within each subject’s complexity metrics maintained a similar shape and range, indicating that these differences between subnuclei were not the product of individual differences between animals and that the sample processing and data acquisition were consistent, supporting the pooling of data between groups (Figure 4A). The inter-individual variability of the fractal dimension values looked different in shape, not only in the non-normal distribution of the data, but also because there was a wider variability below a cumulative probability of 0.5 than above (Figure 4A, right). This was markedly different from the more normally distributed metrics, which showed more consistent patterns of variability across the data.

In terms of the spatial distribution of the values across the dataset, the RN was set apart very clearly from the IO subnuclei with lower curliness and skeleton density values (Figure 4B). This was not as evident in the fractal dimension of the RN ROIs. Generally, in the spatial maps for ROIs within each subnucleus and the RN, the lowest fractal dimension seemed to be located mostly at the margins of the spatial maps (Figure 4B).

Quantitatively, the differences in curliness between the subnuclei and the RN were even more apparent as the anatomical region had a significant effect (Kruskal-Wallis, H = 195.126, DF = 3, p = 4.7669e-42, Figure 4C, left). The MAO and PO had the highest curliness values of 0.284 *±* 0.063 and 0.275 *±* 0.052, respectively (Figure 4C, left, Table 3). The DAO had slightly lower curliness (0.251 *±* 0.060), while it was lowest in the RN (0.197 *±* 0.039, (Figure 4C, left, Table 3). Pairwise comparisons across regions, performed through the calculation of the standard deviation-normalized Wasserstein distance, revealed the largest between the MAO and the DAO at 0.517 SDs (Figure 4C, left). The next largest distance between the PO and the DAO (0.431 SDs), while the difference between the MAO and the PO was the smallest across the comparisons (0.199 SDs, Figure 4C, left). Similarly to the curliness, a significant difference was observed between the skeleton densities of the regions (One-way ANOVA, F = 54.407, p = 6.995e-33, Figure 4C, middle). In terms of effect size across subnuclei, the MAO and DAO had the greatest Wasserstein distance (0.659 SDs, Figure 4C, middle). The distance between PO and DAO was similar at 0.525 SDs, while the MAO-PO distance was the smallest at 0.145 SDs (Figure 4C, middle). In general, this pattern was similar to that of the relationships between the subnuclei in terms of their curliness. Following the pattern of the curliness and skeleton density metrics, fractal dimension was highest in the MAO (1.201 *±* 0.052) and PO (1.197 *±* 0.053, Figure 4C, right, Table 3). Meanwhile, it was lower in the DAO and the RN as their fractal dimension values were 1.163 *±* 0.058 and 1.151 *±* 0.075, respectively (Figure 4C, right). The difference between the fractal dimension of the subnuclei was significantly different (Kruskal-Wallis test, H = 110.0359, DF = 3, p = 1.0780e-23, Figure **??**C). Out of the three metrics described so far, unlike sitting between the values for MAO/PO and RN, the fractal dimension of the DAO was very similar to RN and strikingly different from MAO and PO, as also depicted in the visualization for effect size (Figure 4C, right). More specifically, the effect sizes in the MAO-DAO and PO-DAO comparisons were 0.687 and 0.599, respectively, while the difference between MAO and PO was only 0.095 (Figure 4C, right). This set aside the DAO as more drastically different from the other two subnuclei than in the other metrics.

**Table 3.** Descriptive statistics for the curliness, skeleton density, and fractal dimension across the inferior olive subnuclei and the reticular nucleus in sedentary mice. *n* indicates the number of ROIs. Values are reported as the mean, median, standard deviation (SD), and standard error of the mean (SEM). *N* animals= 9. MAO, medial accessory olive; PO, principal olive; DAO, dorsal accessory olive; RN, reticular nucleus.

| Metric | Subnucleus | *n* ROIs | Mean | Median | SD | SEM |
| --- | --- | --- | --- | --- | --- | --- |
| Curliness | MAO | 478 | 0.284 | 0.284 | 0.063 | 0.003 |
|  | PO | 397 | 0.275 | 0.273 | 0.052 | 0.003 |
|  | DAO | 164 | 0.251 | 0.249 | 0.060 | 0.005 |
|  | RN | 112 | 0.197 | 0.196 | 0.039 | 0.004 |
| Skeleton density | MAO | 478 | 0.784 | 0.790 | 0.188 | 0.009 |
|  | PO | 397 | 0.757 | 0.759 | 0.190 | 0.010 |
|  | DAO | 164 | 0.657 | 0.646 | 0.179 | 0.014 |
|  | RN | 112 | 0.566 | 0.592 | 0.157 | 0.015 |
| Fractal dimension | MAO | 478 | 1.201 | 1.211 | 0.052 | 0.002 |
|  | PO | 397 | 1.197 | 1.207 | 0.053 | 0.003 |
|  | DAO | 164 | 1.163 | 1.166 | 0.058 | 0.005 |
|  | RN | 112 | 1.151 | 1.173 | 0.075 | 0.007 |

#### Differences across the anteroposterior axis

The ROIs from each subnucleus were further divided by anteroposterior position as indicated in Figure 3C. These further classifications resulted in nine distinct groups of ROIs, although, as anticipated, few were classified as posterior DAO and PO (Figure 4D). Furthermore, the lower sampling of the anterior and intermediate DAO is evident. There was no significant difference between these subnuclei with mean curliness values of 0.258 *±* 0.66 and 0.246 *±* 0.057, respectively (Table 4. In the MAO, curliness was highest in the anterior region (0.290 *±* 0.056), then in the posterior (0.286 *±* 0.066), and intermediate regions (0.279 *±* 0.063, Table 4). Although there seems to be a tendency for higher curliness more medial of the IO, especially in the intermediate and posterior MAO in the spatial map, the difference between these subnuclei was not significant (one-way ANOVA, F = 0.915, DF = 2, p = 0.401, Figure 4E). Though not significant, the difference was biggest between the anterior and intermediate MAO as measured by an effect size of 0.216 SDs as compared to the Ant.-Post. and Int.-Post. at 0.161 and 0.140, respectively (Figure 4E). On the other hand, while the anterior PO had a relatively consistent pattern in the spatial distribution of the ROIs, the intermediate PO had higher curliness in the more medial and lateral regions, while the curliness index was lower in the middle across the mediolateral axis (Figure 4D). In the quantification of the comparison within the PO, curliness in the anterior PO was significantly higher than in the intermediate (0.278 *±* 0.052 vs. 0.265 *±* 0.050, Student’s t-test, t = 2.354, DF = 378, p = 0.019, Figure 4E, Table 4.

**Figure 3.**
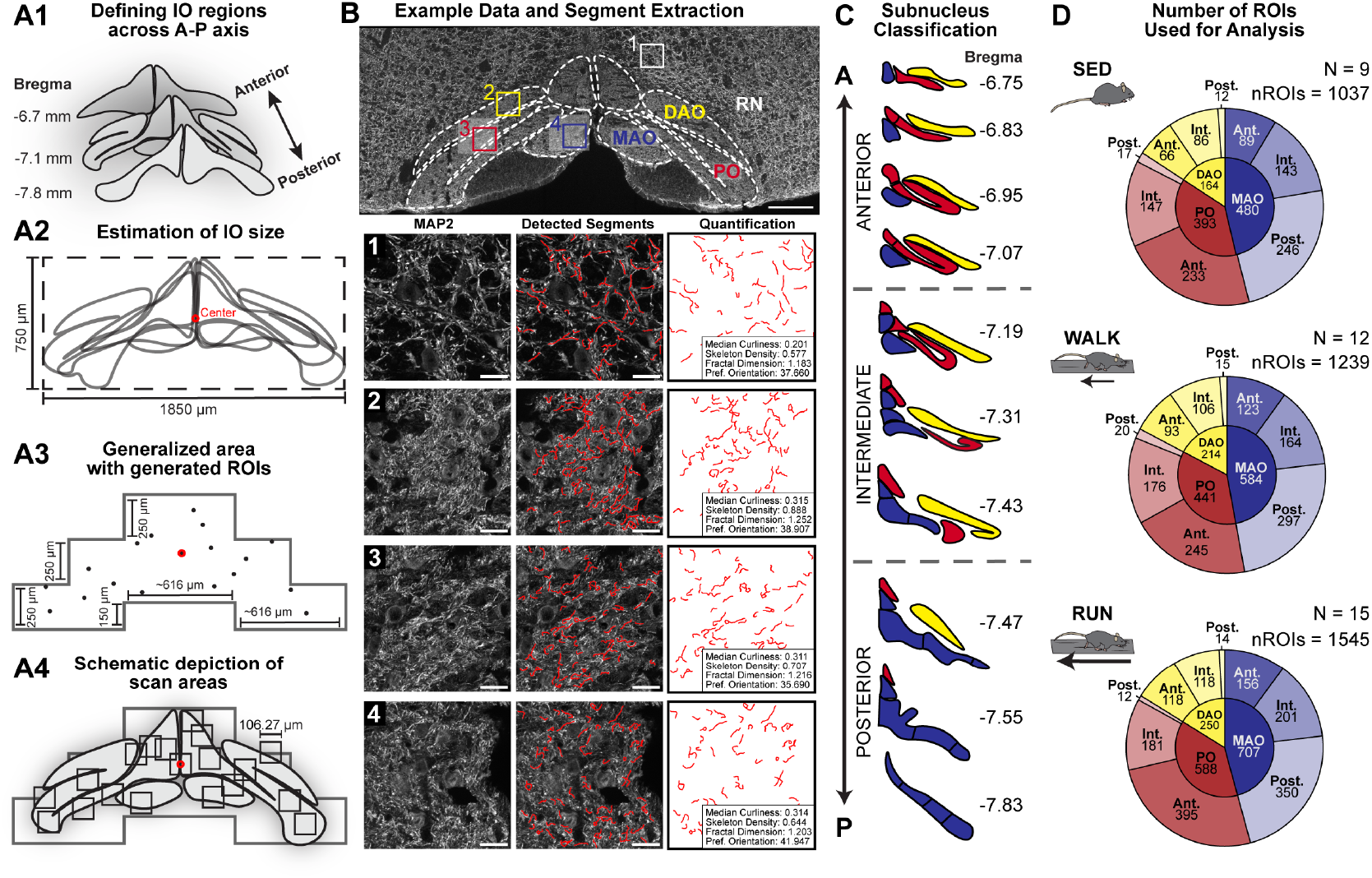
Data acquisition and processing pipeline for high-throughput morphological quantification. (A1) Assessment of the size of coronal IO slices across the A-P axis, (A2) Generalized estimation of the IO area, (A3) subdivision of the area into precise regions and generated ROI locations based on the center coordinate (red), (A4) schematic depiction of ROIs for acquisition setup after generation. (B, Top) Overview image of a coronal slice of the IO stained for MAP2. Scale bar = 200 *µ*m. (B, Bottom) Example maximum intensity projections of MAP2-stained IO neuropil in the 1) RN, 2) DAO, 3) PO, and 4) MAO (left), overlays of the detected segments (middle), and quantifications of the neuropil morphology in each ROI are included (right). Scale bar = 20 *µ*m in 1-4. (C) Subnucleus and A-P location classifications, figure adapted from^59^. (D) The number of ROIs used for the analysis for IO neuropil morphology in the sedentary (top), walking (middle), and running (bottom) groups. N = 9-15/group. DAO, dorsal accessory olive; MAO, medial accessory olive; PO, principal olive; ROI, region of interest; RN, reticular nucleus.

**Table 4.** Descriptive statistics for the curliness, skeleton density, and fractal dimension across the anterior–posterior (A–P) axis of inferior olive subnuclei in sedentary mice. *n* indicates the number of regions of interest (ROIs). Values are reported as the mean, median, standard deviation (SD), and standard error of the mean (SEM). *N* animals = 9. Ant., anterior; DAO, dorsal accessory olive; Int., intermediate; MAO, medial accessory olive; PO, principal olive; Post., posterior.

| Metric | Subnucleus | A–P | *n* ROIs | Mean | Median | SD | SEM |
| --- | --- | --- | --- | --- | --- | --- | --- |
| Curliness | DAO | Ant. | 66 | 0.258 | 0.249 | 0.066 | 0.008 |
|  |  | Int. | 86 | 0.246 | 0.249 | 0.057 | 0.006 |
|  | PO | Ant. | 233 | 0.278 | 0.277 | 0.052 | 0.003 |
|  |  | Int. | 147 | 0.265 | 0.265 | 0.050 | 0.004 |
|  | MAO | Ant. | 89 | 0.290 | 0.290 | 0.056 | 0.006 |
|  |  | Int. | 143 | 0.279 | 0.276 | 0.063 | 0.005 |
|  |  | Post. | 246 | 0.286 | 0.285 | 0.066 | 0.004 |
| Skeleton density | DAO | Ant. | 66 | 0.676 | 0.691 | 0.202 | 0.025 |
|  |  | Int. | 86 | 0.652 | 0.635 | 0.162 | 0.017 |
|  | PO | Ant. | 233 | 0.753 | 0.758 | 0.193 | 0.013 |
|  |  | Int. | 147 | 0.770 | 0.770 | 0.182 | 0.015 |
|  | MAO | Ant. | 89 | 0.745 | 0.733 | 0.181 | 0.019 |
|  |  | Int. | 143 | 0.805 | 0.810 | 0.187 | 0.016 |
|  |  | Post. | 246 | 0.786 | 0.791 | 0.190 | 0.012 |
| Fractal dimension | DAO | Ant. | 66 | 1.158 | 1.165 | 0.067 | 0.008 |
|  |  | Int. | 86 | 1.172 | 1.168 | 0.048 | 0.005 |
|  | PO | Ant. | 233 | 1.197 | 1.207 | 0.054 | 0.004 |
|  |  | Int. | 147 | 1.199 | 1.207 | 0.049 | 0.004 |
|  | MAO | Ant. | 89 | 1.183 | 1.193 | 0.048 | 0.005 |
|  |  | Int. | 143 | 1.206 | 1.216 | 0.051 | 0.004 |
|  |  | Post. | 246 | 1.205 | 1.215 | 0.052 | 0.003 |

In terms of skeleton density of the naive mouse IO neuropil, the anterior region of the DAO had higher skeleton density than the intermediate region (0.676 *±* 0.202 and 0.652 *±* 0.162, respectively, but this difference was not significant (Student’s t-test, t = 0.815, DF = 150, p = 0.416, Table 4). There was very little difference between the mean skeleton densities of the anterior (0.753 *±* 0.193) and intermediate (0.770 *±* 0.182) PO as indicated by a student’s t-test (t = -0.833, DF = 378, p = 0.206) (Figure 4E, Table 4). Unlike the trend of curliness, the anterior MAO had the lowest skeleton density of 0.745 *±* 0.181, while it was highest for the intermediate MAO (0.805 *±* 0.187). The differences between regions were not significant (One-way ANOVA, F = 2.826, DF = 2, p = 0.060, Table 4), but the cumulative probability and the effect sizes show that the posterior region (0.786 *±* 0.190) was much more similar to the intermediate (0.146 SDs), while skeleton density in the anterior MAO stood apart from the two (0.359 and 0.265 SDs from the interior and posterior MAO, respectively (Figure 4E).

**Figure 4.**
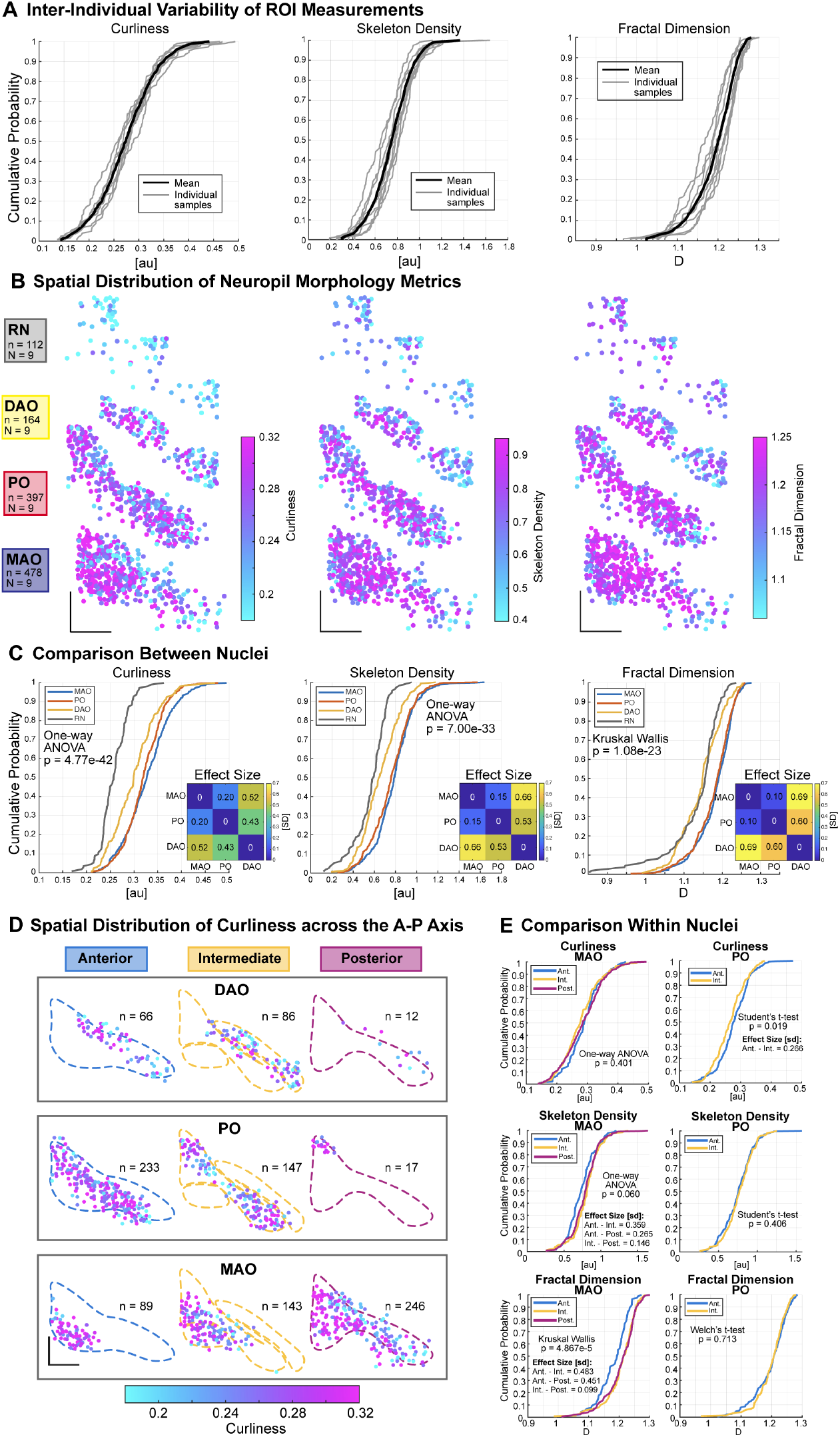
Differences in IO morphology in the naive mouse. (A) CDF of the curliness (left), skeleton density (middle), and fractal dimension (right) from pooled subnuclei acquired from each individual animal (gray) and their mean (black). (B) Spatial distribution of ROIs colored by the curliness, skeleton density, and fractal dimension of segments within each ROI in the RN and each IO subnucleus. Scale bar = 250 *µ*m. (C) CDF of the curliness, skeleton density, fractal dimension, and preferred orientation values within each subnucleus and the RN. Includes the effect sizes measured by the 1D Wasserstein distance representing the differences between each subnucleus normalized to the standard deviation. (D) Spatial distribution of curliness values of each IO subnucleus, subdivided into anterior, intermediate, and posterior regions. (E) Curliness, skeleton density, and fractal dimension in the MAO and PO across the A-P axis. N = 9. DAO, dorsal accessory olive; MAO, medial accessory olive; ROI, region of interest; RN, reticular nucleus.

In the measurement of the fractal dimension of the IO subnucleus, both the intermediate DAO and the PO had a higher fractal dimension. This difference in the DAO was greater as the intermediate region had a fractal dimension of 1.172 *±* 0.048, as compared to that of the anterior region at 1.158 *±* 0.067 (Welch’s t-test, t = -1.441, DF = 113.377, p = 0.152, Table 4). In the PO, the fractal dimension values were generally higher at 1.197 *±* 0.054 and 1.199 *±* 0.049 for the anterior and intermediate regions, but not significantly different (Welch’s t-test, t = -0.368, DF = 336.212, p = 0.713, Figure 4E, Table 4). Meanwhile, in the MAO, the region of the subnucleus had a significant effect on fractal dimension (Kruskal-Wallis test, H = 19.861, DF = 2, p = 4.867e-5, Figure 4E). Similarly to the measurements of skeleton density, the anterior MAO had the lowest fractal dimension (1.183 *±* 0.048, Table 4). Meanwhile, the intermediate and posterior regions of the MAO had not only the highest fractal dimension of this subnucleus, but across subnuclei and regions (1.206 *±* 0.05 for Int. MAO, 1.205 *±* 0.052 for Post. MAO, Table 4). The distance between the fractal dimension values of the anterior MAO to the intermediate and posterior regions was greater compared to the curliness and skeleton density (0.483 from anterior and 0.451 from intermediate, Figure 4E, Table 4).

### Motor experience leads to a localized increase in IO neuropil complexity

Comparison of skeleton metrics in the DAO, PO, and MAO, subdivided along the anteroposterior axis, across the sedentary, walking, and running groups revealed significant differences in the skeleton density and fractal dimension of the intermediate MAO (Figure 5).

**Figure 5.**
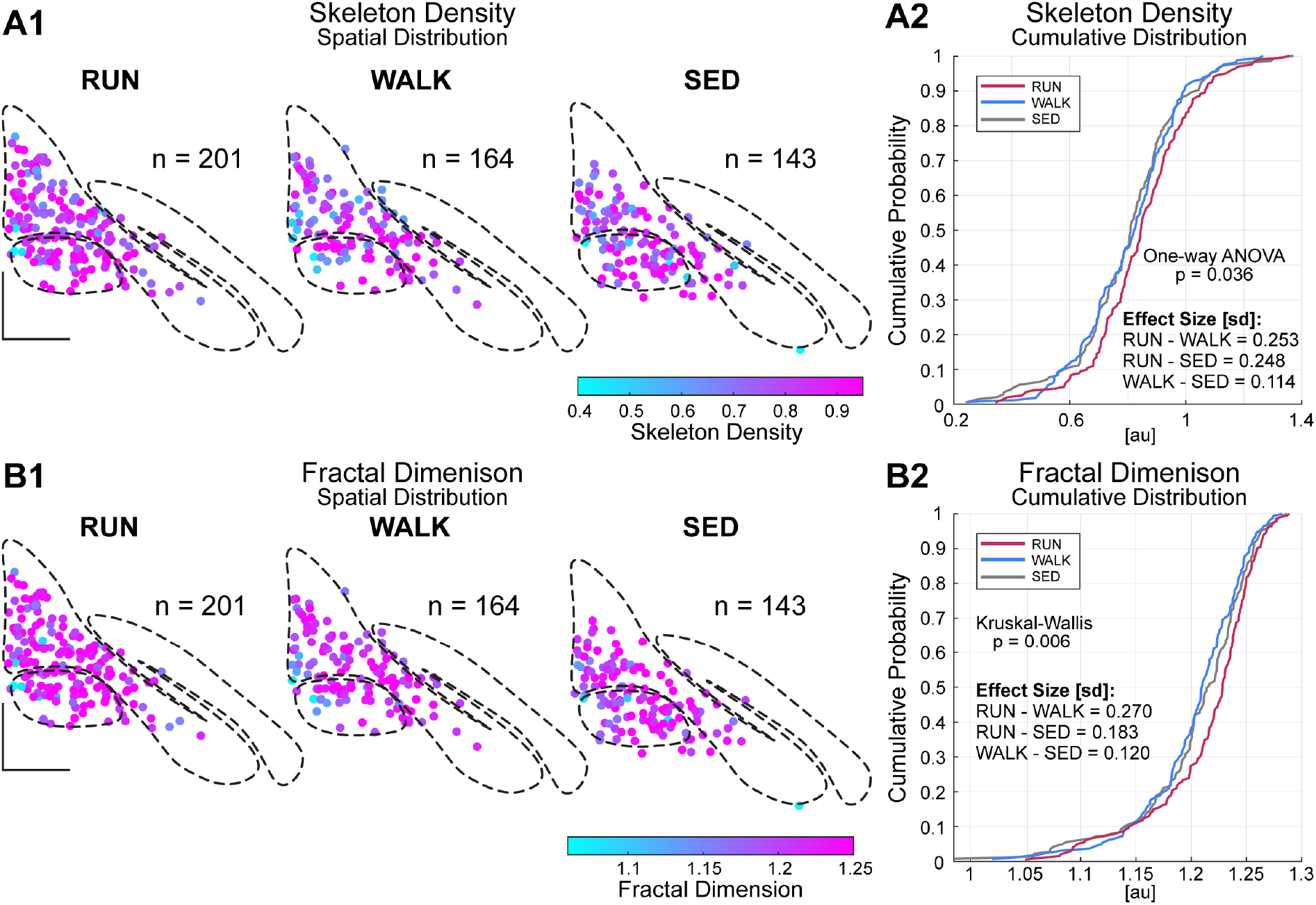
Motor experience significantly increased skeleton density and fractal dimension of the intermediate MAO. (A1) Spatial distribution plots of skeleton density in the intermediate MAO in the running, walking, and sedentary groups. (A2) Cumulative distribution function of skeleton density, including effect size as measured by the SD-normalized Wasserstein distance. (B1) Spatial distribution plots of fractal dimension in the intermediate MAO in the running, walking, and sedentary groups. (B2) Cumulative distribution function of fractal dimension. N = 9-15/group. Scale bar = 250 *µ*m.

A significant difference was found in the skeleton density of the intermediate MAO across the behavioral groups (One-way ANOVA, DF = 2, F = 3.354, P = 0.036, Figure 5A2). In observing the spatial distribution of the ROIs and their skeleton density values, this effect is not immediately obvious (Figure 5A1). Nevertheless, the sedentary and walking groups were close in skeleton density with values of 0.805 *±* 0.187 and 0.809 *±* 0.169, respectively, but the skeleton density in the running group was higher at 0.848 *±* 0.174 (Table5). This is about 5.342 % higher than the sedentary group and 4.821 % higher than the walking group. This was also reflected in the effect sizes (Figure 5A2). The effect of group on fractal dimension was also not immediately obvious by eye, nor did any hotspots stand out in the spatial distribution of the metric, and as the varying number of ROIs from each group also affects the visualization of these maps, color-coded by the metric (Figure 5B1). However, the fractal dimension in the intermediate MAO was also significantly different across behavioral groups (Kruskal-Wallis, H = 10.3474, DF = 2, p = 0.006, Figure 5B2).

Aligned with the skeleton density results, the running group’s intermediate MAO had the highest fractal dimension (1.215 *±* 0.047, Table 5). The fractal dimension of the sedentary and walking groups was similarly closer to one another than to the running group, but the sedentary int. MAO had a higher fractal dimension at 1.206 *±* 0.051 than that of the walking group at 1.204 *±* 0.046 (Table 5). This was a 0.914 % increase from the walking group and a 0.746 % increase from the sedentary group, while the difference between the walking and sedentary groups was only 0.166 %. Therefore, the greatest difference was between the running and walking group with a 0.270-SD effect size as compared to the running-sedentary effect size of 0.183 (Figure 5B2).

**Table 5.** Descriptive statistics for the skeleton density and fractal dimension in the intermediate medial accessory olive (MAO) across sedentary, walking, and running groups. *N* indicates the number of animals and *n* indicates the number of ROIs. Values are reported as the mean, median, standard deviation (SD), and standard error of the mean (SEM).

| Metric | Group | *N* animals | *n* ROIs | Mean | Median | SD | SEM |
| --- | --- | --- | --- | --- | --- | --- | --- |
| Skeleton density | SED | 9 | 143 | 0.805 | 0.810 | 0.187 | 0.016 |
|  | WALK | 12 | 164 | 0.809 | 0.824 | 0.169 | 0.013 |
|  | RUN | 15 | 201 | 0.848 | 0.846 | 0.174 | 0.012 |
| Fractal dimension | SED | 9 | 143 | 1.206 | 1.216 | 0.051 | 0.004 |
|  | WALK | 12 | 164 | 1.204 | 1.210 | 0.046 | 0.004 |
|  | RUN | 15 | 201 | 1.215 | 1.228 | 0.047 | 0.003 |

**Table 6.** Sample sizes and punctum counts used for the Cx36 subnucleus analysis. An ROI is one acquired scene. Punctum counts are the sums across included ROIs, and mean puncta per ROI are reported as mean *±* SD across ROIs.

| Subnucleus | *n* animals | | *n* slices | | *n* ROIs | | *n* puncta | | Puncta per ROI, mean ± SD | |
| --- | --- | --- | --- | --- | --- | --- | --- | --- | --- | --- |
|  | SED | RUN | SED | RUN | SED | RUN | SED | RUN | SED | RUN |
| MAO | 3 | 3 | 9 | 10 | 18 | 20 | 90,504 | 131,842 | 5,028 ± 1,140 | 6,592 ± 1,965 |
| mPO | 3 | 3 | 10 | 10 | 20 | 20 | 110,860 | 131,227 | 5,543 ± 1,812 | 6,561 ± 1,821 |
| vlPO | 3 | 3 | 10 | 8 | 16 | 14 | 97,059 | 90,290 | 6,066 ± 2,150 | 6,449 ± 2,212 |
| RN | 3 | 3 | 10 | 10 | 10 | 10 | 57,443 | 64,748 | 5,744 ± 1,276 | 6,475 ± 2,224 |

### Localized changes observed in Cx36 signal as a result of motor experience

If the dendritic neuropil indeed grows in complexity in response to motor experience, how could it lead to changes in olivary activity? As the only known way in which IO neurons communicate with each other involves electrical coupling via Cx36- formed gap junctions, we proceeded to examine whether evidence for changes in gap junction density could be found. For this purpose, medullary sections from three randomly selected animals from the sedentary and running groups were processed for immunohistological demonstration of Cx36, and 63x confocal image stacks were acquired from MAO, medial PO, and ventrolateral PO localized at the intermediate anteroposterior levels (Figure 6A, B). In total, 38, 40 and 30 image stacks were included in the analysis at the aforementioned IO locations after quality control exclusions, with an additional 20 stacks acquired at the nearby region of reticular nucleus (see Figure 6C for locations in sedentary and running groups). For each image stack, fluorescent puncta locations were extracted, and the 3D density of puncta was calculated in 5-um neighborhoods (Figure 6d).

**Figure 6.**
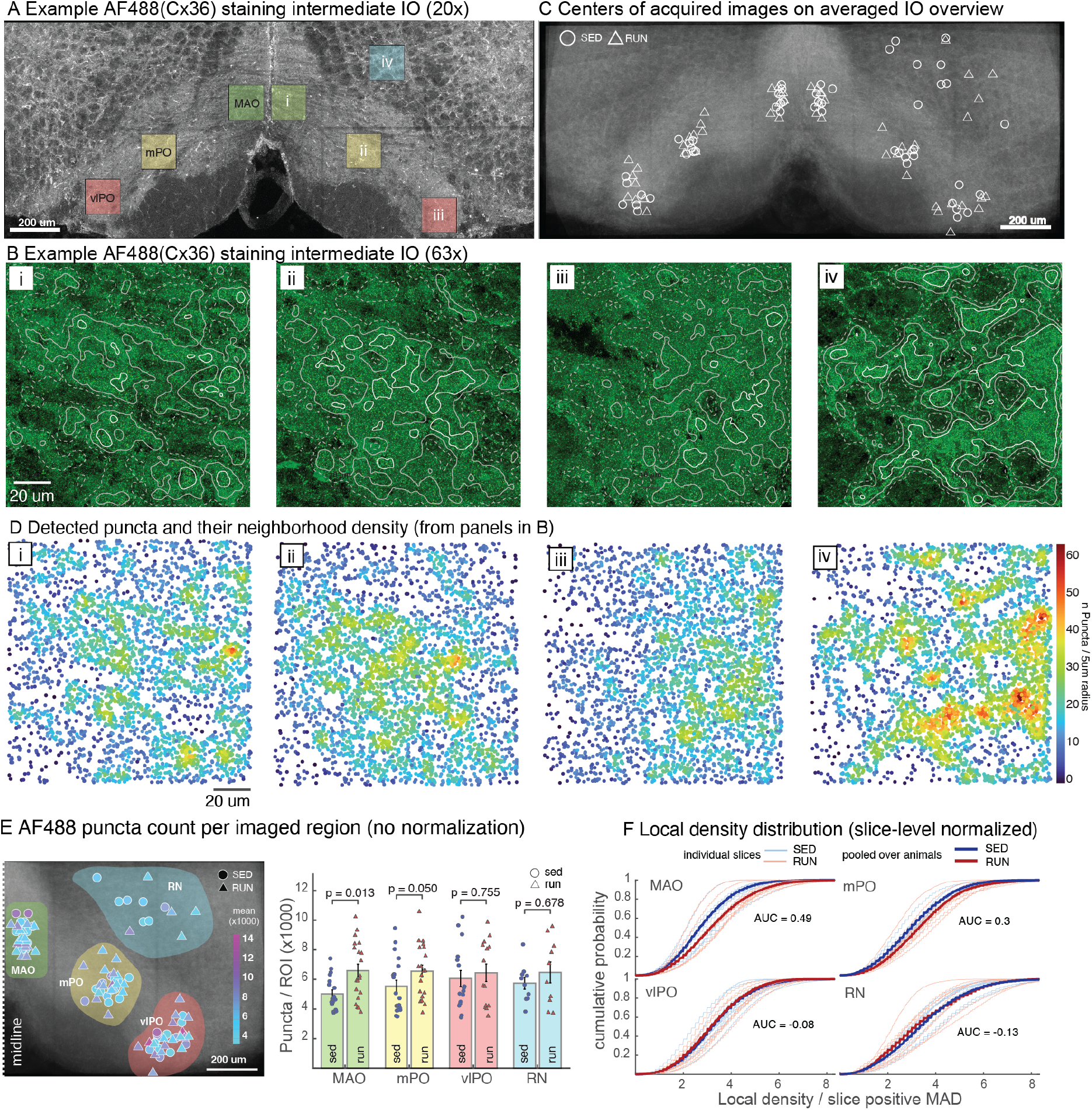
Motor experience results in localized changes in Cx36 puncta. Cx36 detection in animals with and without running experience. A, example 20x overview standard-deviation projection image of an inferior olive processed for immunohistological detection of Cx36. Square regions indicate locations of high-resolution image stacks acquired for this slice. B, standard deviation projection images for regions indicated i-iv in panel A. C, centers of image locations shown overlaid on an average of IO slices used in this part of the work. Locations for sedentary and run groups (indicated by circles and triangles, respectively) are intermingled. Note that locations from 2 slices are not shown in this panel, as the sample was too distorted for automatic positioning. D, detected AF488 puncta obtained from the panels shown in B. Color code indicates the local density, defined as the number of puncta found within a 5-micrometer radius. E, left: number of puncta detected within an image shown overlaid on the averaged IO image, with left and right sides mirrored and superimposed. Color-shaded regions indicate location classification; darker colors of markers indicate images with higher counts. Right: AF488 puncta counts per image (markers) are significantly higher in the run group for MAO locations. Bars denote averages *±* standard deviation. p-values denote wilcoxon rank sum. F, distributions of local density metrics across all acquisitions (thin lines) and their averages (thick lines). The density values for each image are normalized by slice-level median absolute deviation (MAD). Area Under Curve (AUC) values show a considerable increase in density values for MAO and mPO. Abbreviations: MAO, medial accessory olive; mPO, medial principal olive; vlPO, ventrolateral principal olive; RN, reticular nucleus.

In this dataset comprising 10 slices in both groups and up to 20 image stacks from sed and run groups for each area of interest (see table), were analyzed. Intriguingly, a significant increase was seen in the number of puncta detected in each image in the MAO region (mean n of puncta per image: 5028+-1140 and 6592 +- 1965 for sedentary and runners, p = 0.013, Wilcoxon rank-sum; Figure 6E).

To ameliorate the unavoidable effects of slice-level sample quality and staining variability, local Cx36 punctum-density distributions were compared using cumulative distribution functions (Figure 6F). For each animal, local density values were normalized to the median absolute deviation of positive density values within each slice, and punctum density values for a subnuclear location in one animal were pooled. Again, samples obtained from the running group showed rightward shifts toward higher local densities in MAO and mPO (area under curve differences: 0.49 and 0.30), but little or no corresponding shift in vlPO or RN (-0.08 and -0.13). Therefore, it is plausible that in the regions of the IO where an increase of neuropil complexity was observed (especially intermediate MAO), gap junction density is also increasing.

## Discussion

In this work, we provide (to our knowledge) the first evidence for experience-induced plasticity in the inferior olive. Below, we will elaborate on the details and limitations of these first observations. However, even with the caveats, the results indicate that the olive is not a hard-wired relay station and call for further studies examining the physiological and molecular mechanisms underlying olivary plasticity.

### The neuropil is not uniform across IO subnuclei

A classic notion regarding the olivo-cerebellar systems has been that of homogeneity: that it is comprised of virtually identical and repeating micromodules with uniform structure, and hence any differences in outputs of these modules are driven by processes upstream to them. This elegant and convenient model has been gradually replaced as studies describing anatomical and physiological differences among the cerebellar cortical and nuclear regions (^4,39–42^) have been published. However, despite several attempts (^9,43–46^) only weak structural or physiological differences have been described for the inferior olive with the exception of the cerebellar flocculus-targeting dorsal cap of Kooy. Notably, the studies that have been technically limited to a relatively small number of indvidual olivary neurons may have missed subtle or gradual differences between regions.

In the present work, we sought to sample the IO subnuclei densely enough to overcome the barrier of single-cell variability. As obtaining a large number of single-cell measurements - such as patch-filled morphologies - from animals in three experimental groups at a defined time point was not technically feasible, we chose to conduct the morphological examination of the olive in a neuronal identity-agnostic manner. Building on the notion that computation in a network with dendro-dendritic communication and dendrite-emerging axons^47,48^ is not necessarily defined by the soma, we quantified the shapes of short, unbranched segments extracted from a large number of randomly-positioned image stacks acquired from MAP2-stained IO slices from animals that had not experienced any motor training.

Curiously, when pooled across all measurement locations, the neuropil in the medial accessory olive region was found to be more complex than that of the other nuclei, and the dorsal accerrosy olive least complex. If these differences reflect a lower incidence of dendritic appositions and electrical coupling in the DAO, it would be directly in line with our earlier finding that the coactivation levels of dorsal accessory nuclei are lower than those of the principal olive^38^. Notably, while this complexity order of MAO - PO - DAO seems to contrast previous reports describing lower-complexity (“Type I”) neurons being found in the MAO^8^, our earlier work^9^ found that curly morphologies are not less common in the MAO than the PO. It might be that the curved, sheet-like shape of the PO is not conducive to preservation of long, straight dendrites in coronal sections, and hence they would be underrepresented in single-cell-labeling observations from the PO.

### Localized increases in neuropil complexity after motor experience

In addition to the differences among the subnuclei in naive animals, we found consistent changes in the measurements when comparing animals with different motor experiences. Specifically, we saw a clear increase in metrics that describe neuropil complexity in the intermediate medial accessory olive in animals that had completed the 2-week treadmill running challenge, contrasting the measurements in the sedentary and walking group animals. The complexity-enhanced IO region is considered to be involved in mechanisms underlying balance (^49^), and the putative increase of its neuropil’s complexity - dendritic or axonal - would suggest that the development of skilled high-speed treadmill running requires improving balance. While our balance-beam testing paradigm did not reveal major performance improvements, it is possible that on a more challenging test (with a thinner and/or longer beam), the trends for faster traverse speed and less stopping in the running group would have become significant.

### Evidence for changes in gap junction coupling after motor experience

Due to limited time and resources, we had to restrict our examination of Cx36 labeling to a small number of animals (3) and locations within the IO. Nevertheless, using two metrics - absolute number of AF488-labeled puncta per each acquired image stack, and distribution of localized density measurements normalized by robust slice-level median - the region in the interposed MAO showed a clear and significant increase in labeling. Intriguingly, we also saw an increase localized in the medial part of the ventral fold of the PO linked to the cerebellar zone D1 (^50^) related to visual control of voluntary, skilled locomotion(^51,52^). In contrast, no changes were seen in the ventrolateral bend region of the olive, projecting to the cerebellar Crus 1/2 region linked to flexible and predictive behavior. Indeed, as the treadmill running task was designed to be as predictable and uniform as possible, changes in this region were not expected.

### Methodological considerations and limitations of the study

Our methodological approach allows measuring morphological features of the IO neuropil in a high-throughput manner across all the IO subnuclei. While efficient and specific to neuronal components of the neuropil (^53,54^), the work has the obvious limitation that we can not assign observed changes to distinct cells or cell types. Furthermore, without reconstruction of individual dendrites (as was done in^9^), it is possible that the segment-level measurements obscure changes occurring at the scale of full dendrites. Hence, an increase in “dendritic complexity” could either result from dendrite tips becoming more curly or from dendrites growing more branches. Nevertheless, if our admittedly crude measures obtained from relatively large regions of interest reveal differences, it is likely that more targeted studies (e.g. by means of sparse viral labeling,^55^) would uncover more dramatic changes.

Along the same lines, our results, including the observation of increased Cx36 labeling in the regions where neuropil complexity was increased, can not be taken as proof of increased electrical coupling between IO neurons. Even if the number and density of GJs between two neurons would increase, the effective coupling would only be enhanced if the GJ density as well as the proportion of connexin channels in open state increased with respect to the coupled dendritic volumes. The definitive experimental proof for upmodulation of electrical coupling after motor experience will require targeted paired patch-clamp or dye-coupling experiments, both of which are challenging to conduct in living animals.

### Significance for olivo-cerebellar function and learning

Despite the methodological shortcomings, the results of this work are novel and establish the notion that the olivary dendritic neuropil is modifiable in healthy adult animals, and lay the foundations to consider the IO as an active, plastic node in the network, instead of a hard-wired source of the CF. Curling or the increase in density within the olivary neuropil could therefore be a long-term form of plasticity that would not only up- or downregulate coupling strength^56,57^ at the existing gap junction sites but also modify the landscape of possible contact points. How could such changes be predicted to affect complex spike activity in the cerebellar cortex?

Increasing not only single-site coupling strength but also the number of GJ sites could obviously lead to changes in levels of co-activation between IO neurons and, thereby, of cerebellar complex spikes. Furthermore, as afferent inputs might be more broadly shared among strong-coupled neurons, the “receptive domains” of individual IO neurons that define the pre-olivary afferent sources capable of driving spike generation may broaden, allowing single neurons to respond to more diverse inputs. On the other hand, increased density of gap junctions is expected to result in elevated leak currents, lower input resistance, possibly affecting single-cell excitability as well as time constants related to post-spike refractory period.

Thus, while it is tempting to predict that an increase in electrical coupling in the IO resulting from motor experiences or learning would lead to enhanced levels of co-activation of CSs, more work, including single-cell-level analysis of the physiological and morphological changes is needed to complete the picture.

## Methods

### Animal Subjects

Male wild-type C57BL/6J mice (CLEA Japan, Inc., Japan) aged 8-10 weeks (postnatal P56-70), were housed at a reverse cycle (12 hours dark, 12 hours light), equipped with a house, a tube, and bedding, and had access to food and water *ad libitum*. A wheel was not provided inside the cage. Experiments were conducted in accordance with the relevant guidelines and regulations set by the Okinawa Institute of Science and Technology Animal Care and Use Committee (ACUC), an Association for Assessment and Accreditation of Laboratory Animal Care (AAALAC International)-accredited facility under protocol number 2023-053. This study was reviewed and approved by the Okinawa Institute of Science and Technology Animal Care and Use Committee (ACUC).

### Pre-experiment Handling and Training

Mice were handled one week after arrival to the facility for approximately five days and consisted of acclimation to the experimental room and experimenter by 1) taking them out from the cage with a tube, 2) gently pulling by the tail base onto the forearm or outer part of the hand with constant support of the paws, 3) scooping with one hand holding the tail gently, and 4) scooping without holding the tail. All behavioral recordings were performed no earlier than two weeks after arrival at the facility, and the animals were brought to the experimental rooms at least one hour before starting training or recording on each day to allow adequate acclimation.

### Experiment

The behavioral experiments consisted of treadmill training for 14 consecutive days. Body weight was measured each day using a bench scale (AXEL Corporation, Japan) before training or recordings. Balance beam and grip strength tests were performed at three time points during the 14-day training period: days 7 and 14, and a baseline recording (day 0). Animals were divided into three groups based on the training or lack thereof across the 14 days. The behavioral groups based on treadmill training were the running (N = 18), walking (N = 12), and sedentary (N = 12) groups. The target treadmill speeds were 30 and 15 m/min for the running and walking groups, respectively.

#### Treadmill

The mice were trained on a single-lane mouse treadmill (Maze Engineers, USA). No shock grid or plexiglass cover was used. The day before beginning treadmill training, animals were exposed to the treadmill environment and surroundings by placing them onto a 30 by 30 cm plastic grid positioned on top of the treadmill, without walls, to allow the mouse to acclimate to the height and lighting of the treadmill setting. The grid was also used specifically to avoid exposure to a stationary treadmill. The mouse was prevented from leaving the arena by taking it back by the tail if it attempted to climb out. The mouse was kept in the arena for one minute with the treadmill on underneath the grid to allow habituation to the sound of a running treadmill. On each experimental day, the mice were brought into the experimental room an hour before the start of the experiment to allow for acclimation to the environment. During training, verbal encouragement was given by the experimenter. Furthermore, the experimenter used their hands to encourage running by placing a hand at the front of the treadmill or lightly tapping the mouse from behind, making sure that the mouse was not being pushed or that it was relying on the hands to continue running. In the rare case that the mouse refused to run, it was not forced to do so. In addition to the changes made to treadmill training and criteria, the timelines for the relevant behavioral paradigms were slightly different. The overall timeline of these behavioral tests is shown in the figure below.

#### Balance Beam Test

The mice were introduced to the balance beam setup two days before the baseline recording and were trained to cross the beam. One day prior to recordings on days 7 and 14 of treadmill, the mice were reacclimated to the setup, and each mouse crossed the beam once. This was repeated if the mouse fell off or refused to cross. On the recording days, three trials of crossing a 70 cm-long and 7 mm-wide steel balance beam were recorded using a single flanking camera on the left side of the mouse at 160 fps (DMK 37BUX273, The Imaging Source, USA). A trial was not considered if the mouse fell off or flipped over on the balance beam and was repeated. The trials were averaged for each day.

#### Grip Strength Test

Forelimb and four-limb grip strength were measured using a grip strength meter (Bioseb, France) with a grid grip by gently pulling the mouse away from the apparatus horizontally by the base of the tail. In the forelimb grip strength test, the mouse was only allowed to grasp the top of the grid with its forepaws. Grip strength was normalized to individual body weight on the respective recording day.

### Brain Slice Preparation

The animals were transcardially perfused with room-temperature 4% paraformaldehyde (PFA, Electron Microscopy Sciences, USA) in phosphate-buffered saline (PBS, Gibco, Thermo Fisher Scientific, USA) using a peristaltic pump (Perista AC-2110, ATTO, Japan) after subcutaneous injection of 100 *µ*L (100 mg/mL) phenobarbital, and confirmation of deep anesthesia. The brains were extracted and stored in 4 % PFA for 2-4 hours for slices stained for MAP2. Slices stained for Cx36 were stored directly in PBS without further postfixation. The tissue was cryoprotected using stepwise concentrations of sucrose (Nacalai Tesque Inc., Japan) in PBS. The tissue was incubated in 10 and 20 % for 3 hours at room temperature, and in 30 % overnight at 4 °C. The tissue was washed using PBS and embedded in Tissue-Tek® O.C.T. Compound (Sakura Finetek USA Inc., USA) inside Peel-A-Way™ embedding molds (Sigma-Aldrich, USA), and placed onto thermal beads cooled down to approximately -80 °C to solidify. The brains were coronally sectioned at 50 *µ*m using a cryostat (CM1950, Leica Biosystems, Germany) and stored in PBS. Slices were mounted onto glass microscopy slides (1338 Globe Scientific Inc., USA), with VECTASHIELD antifade mounting medium (H-1000, Vector Laboratories, USA), covered with No. 1.5H cover glass (Marienfeld, Germany), and sealed using CoverGrip^TM^ Coverslip Sealant (23005, Biotium, USA). Exactly nine slices on one slide were mounted from each animal.

### Immunohistochemistry

The tissue was rinsed 4 times with PBS for 10 minutes per wash. The slices were incubated in blocking and permeabilization buffer (10% bovine serum albumin (Sigma-Aldrich, USA), 1% normal goat serum (ab7481, Abcam, United Kingdom), 0.4 % Triton^TM^ X-100 (Sigma-Aldrich, USA) in PBS) for 60 minutes at room temperature. For MAP2 labeling, the slices were incubated with the primary chicken polyclonal anti-MAP2 antibody (ab5392, Abcam, United Kingdom) diluted 1:1000 in PBS containing 2% BSA and 0.4 % Triton^TM^ X-100 at 4 °C with gentle shaking (80 rpm) for 48 hours. This was followed by 4 washes of 10 minutes with and incubation with the secondary goat anti-chicken IgY antibody, Alexa Fluor 488 (A-11039, Thermo Fisher Scientific, USA), diluted 1:2000 in PBS containing 2% BSA and 0.4 % Triton^TM^ X-100 for 3 hours at room temperature, on a shaker (80 rpm), protected from light, followed by 4 washes of 10 minutes each. For Cx36 labeling, slices were incubated with the primary mouse monoclonal antibody (1E5H5, Thermo Fisher Scientific, USA) diluted 1:200 in PBS containing 5% BSA and 0.4 % Triton^TM^ X-100 at 4 °C with gentle shaking for 72 hours. After 4 washes of 10 minutes, the slices were incubated with the polyclonal secondary goat anti-mouse antibody, Alexa Fluor 488 (A-11001, Thermo Fisher Scientific, USA) diluted 1:2000 in PBS containing 5% BSA and 0.4 % Triton^TM^ X-100 for 3 hours at room temperature. Slices were washed 4 times for 10 minutes each.

### Image Acquisition

For MAP2-stained slices, 16-bit images of the immunohistochemically labeled brainstem slices were acquired using a Zeiss LSM 880 confocal laser scanning microscope (Carl Zeiss Microscopy GmbH, Germany). Overview images of the brainstem were acquired using a 10x objective (Plan-Apochromat 10x, NA = 0.5; Carl Zeiss Microscopy, Germany), with average 2, speed 9, zoom 1.1, 0.76 *µ*m per pixel, laser power 2 %, master gain 600, digital gain 1, pinhole 1 AU (28.8), 8-10 z-steps with a z-increment of 5 *µ*m, 7x4 tiling, 6867x3951 image size. Additionally, 20x overview images were acquired of the IO using a 20x objective (Plan-Apochromat 20x, NA = 0.8; Carl Zeiss Microscopy, Germany) with average 2, speed 9, zoom 1, 0.42 *µ*m per pixel, laser power 1.5 %, master gain 600, digital gain 1, pinhole 1 AU, 10 z-steps with 1 *µ*m z-increment, 5x2 tiling, 4920x2005 image size. Finally, high-magnification ROI images were acquired using a 40x objective (Plan-Apochromat 40x Oil DIC M27, NA = 1.4; Zeiss Immersol oil; Carl Zeiss Microscopy, Germany) with average 2, speed 6 (pixel dwell 2.05 *mu*s), zoom 2, 0.10 *µ*m per pixel, laser power 1 %, master gain 600, digital gain 1, pinhole 1 AU (28.8 *µ*m), 30 z-steps with 0.2 *µ*m z-increment, 1024x1024 image size. The excitation and detection wavelengths were 488 nm and 481–555, respectively. High-magnification acquisition of Cx36-stained brainstem slices were acquired using a 63x objective (Plan-Apochromat 63x Oil DIC M27, NA = 1.4; Zeiss Immersol oil; Carl Zeiss Microscopy, Germany) with average 2, speed 6 (pixel dwell 2.05 *mu*s), zoom 1, 0.13 *µ*m per pixel, laser power 1.5 %, master gain 600, digital gain 1, pinhole 1 AU (49.1 *µ*m), 51 z-steps with 0.1 *µ*m z-increment, 1024x1024 image size. The excitation wavelength was 488 and the detection wavelength was 481–555 nm.

### Data Processing and Analysis

#### Balance Beam Performance Analysis

The video size was 1280x376 acquired at 160 fps. The raw videos (.avi) were preprocessed by converting them to a nearly lossless .mp4 file by calling FFMPEG^58^ (version 8.0). The analysis pipeline steps were (1) bar detection, (2) mouse tracking, and (3) the detection of metrics such as the number of slips, mean speed, and the number of stops. The bar was detected in the video based on the location of pieces of dark tape at the ends of the bar, and the mouse was tracked using thresholding and creating a binary mask for each frame. The calculation of the time it took the mouse to traverse the beam, the mean speed of the cross, and the detection of stops were based on the centroid, as well as user-set parameters such as the threshold for mouse locomotion in pixels per second. Furthermore, the determined stops were used to calculate the mean locomotion speed (distinct from mean speed), which excluded detected stops from the speed calculation. The distance from the middle of the bar and the y-coordinate of the mouse centroid were used to calculate a mean postural height metric of the mouse as it crossed. Slips were detected by measuring changes in pixel intensity in a designated area under the bar. Slips were only considered when they occurred under the body of the mouse. The formal description of the algorithm can be found in the BEAMCROSS GitHub page.

#### Preprocessing and Skeletonization of High-resolution MAP2-Stained ROIs

Skeletonization of MAP2-stained images was performed by computing a z-projection (SD) of the 6-*µ*m thick ROIs. Then, extreme-intensity-value outliers were removed using contrast (histogram) clipping and stretching, setting the minimum and maximum intensity values to the 1-99.5 percentiles of the original z-projection’s full intensity range. Further steps were taken to dampen outliers, often in the form of very bright signals due to dirt or artifacts in the scans, by detecting unusually bright pixel regions using a robust, log-space z-score. Only large, connected, bright areas were retained, and the unusually bright pixels were replaced with pixels of a typical background intensity taken from the histogram of the remaining image. Next, a “likelihood map” of the dendritic fibers was computed using a Frangi-based vesselness filter, followed by binary masking. Skeletonization was performed on this likelihood map, producing single-pixel wide skeletons of the segments. Branches in the skeletons were eliminated by removing pixels at the branch points. This process depended on user-set parameters: minimum segment length, a range of acceptable segment counts within the ROI, and the minimum contrast threshold between the minimum and maximum intensity values in the average intensity plotted along the z-axis of the raw scan before z-projection. These parameters allowed automatic exclusion of out-of-focus and off-target ROIs. If branches within each ROI were shorter than 50 pixels (approximately 5.19 *µ*m), they were excluded. The minimum and maximum numbers of segments per ROI were 10 and 200, respectively. If outside of this range, the ROIs were excluded from the dataset. The contrast threshold between the brightest and darkest points of the image was 200 to exclude ROIs that were out of focus in the z-plane. These parameters were kept consistent across datasets.

#### Anatomical Classification of IO Subregions

After acquisition of high-resolution ROIs of the MAP2-stained IO neuropil, and before analysis of the morphology, each slice was manually assigned tags indicating its position on the anteroposterior axis of the IO. The classifications were anterior, intermediate, and posterior and were based on published descriptions of IO structure. While the assignment was done manually, it was supported by a GUI to provide anteroposterior and subnucleus tags for each ROI in an appropriate format for integration into the later analysis algorithm. Since slices were sampled from across the A-P axis of the IO in these experiments, the classifications were made across the axis. The approximate threshold for differentiating anterior and intermediate IO was between Bregma -7.19 and -7.07. This threshold for intermediate versus posterior was between Bregma -7.47 and -7.43. This was decided on because each of these regions had unique shapes that were identifiable from the 20x overview scans. The definition of the PO in this work is comprised of the major principal nucleus, the arcuate subnucleus, the IOK, the IOVL, and the IODM. The definition of MAO encompasses the major MAO nucleus, the IOBe, the IOA, the IOB, and the IOC, while the DAO is composed of the major DAO and the IODdf.

#### Neuropil Complexity Measurements

Data processing was performed on SD z-projections of the high-resolution ROIs. Extreme intensity value outliers were removed using contrast (histogram) clipping and stretching. This set the minimum and maximum intensity values as the 1 - 99.5 percentiles of the full intensity range of the original z-projection. Further steps were taken to dampen outliers by detecting unusually bright pixel regions using a robust, log-space z-score. Only large, connected, bright areas were retained, and the unusually bright pixels were replaced with pixels of a typical background intensity taken from the histogram of the remaining image. A “likelihood map” of the dendritic fibers was computed using a Frangi-based vesselness filter, followed by binary masking. Skeletonization was performed on this likelihood map, producing single-pixel wide skeletons of the segments. Branches in the skeletons were eliminated by removing pixels at the branch points.

In the quantification step, some user-set parameters were set. These were a minimum segment length of 50 pixels (approximately 5.19 *µ*m), a segment count range of 10-200 per ROI, and an intensity contrast threshold value across the z-axis of 200.

Curliness of each ROI was calculated as the ratio of each segment’s total length to the Euclidean distance between its endpoints, and the median of all segments within said ROI was computed. Skeleton density was calculated by dividing the total length of the skeleton by the total detected labeled area. Fractal dimension was calculated using the box counting method.

#### Quantification of Connexin-36 Density

AF488 puncta, putatively indicating the presence of Cx36-formed gap junctions, were quantified from 63x confocal image stacks (xy resolution: 0.13 um; z-step 0.1 um). The detection was done using MATLAB code (the GitHub repository will be publicly available by the final submission).

Before detection, large bright artifacts defined as the 98th percentile of all voxel intensities with areas larger than 1 um in diameter were identified in each image and inpainted with local median values.

Puncta were detected using a multiscale three-dimensional Laplacian-of-Gaussian (LoG) detector. LoG responses were calculated at physical Gaussian scales of

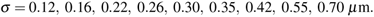

For each scale, Gaussian smoothing was adjusted according to the physical voxel size, and the LoG response was multiplied by *−σ* ^2^ so that compact bright structures produced positive responses. The response at each scale was robustly normalized using the median and median absolute deviation:

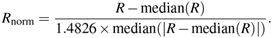

For each voxel, the maximum normalized response across scales was retained.

Candidate puncta were defined as 26-connected regional maxima in the multiscale response image. Candidates were required to exceed a normalized LoG response threshold of 20. When a regional maximum formed a multi-voxel plateau, only the voxel with the strongest response was retained.

Candidate puncta were then validated by measuring their local XY size in the original image plane. Around each seed, a local image patch was smoothed with a 0.5-pixel Gaussian filter. The local background was estimated from an annulus extending from 1.0 to 1.5 *µ*m from the seed. The local peak intensity was searched within one pixel of the LoG seed. Candidates were rejected if the local peak did not exceed the local background or if fewer than 10 pixels were available for background estimation.

For each remaining candidate, a half-prominence threshold was defined as

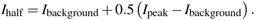

The foreground component connected to the local peak was segmented at this threshold using 8-connected two-dimensional connectivity. Component area was converted to an equivalent circular XY diameter:

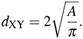

Candidates were rejected if the connected component touched the local patch boundary or if *d*_XY_ *>* 1.0 *µ*m. Size, shape or fluorescence intensity of the puncta was not examined in the analysis.

Local puncta density was calculated for each accepted punctum by counting neighboring puncta within a three-dimensional sphere of radius 5 *µ*m. The punctum itself was excluded from the neighbor count. Local density was defined as

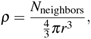

where *r* = 5 *µ*m. Densities are reported as puncta per cubic micrometer. No correction was applied for the truncation of the neighborhood sphere at stack boundaries, nor for possible tissue compression in the z-dimension.

For spatial visualization (Figure X), accepted puncta were projected into the XY plane. Projected density maps were calculated by counting puncta within a 5 *µ*m radius disk around each image pixel and dividing by disk area, yielding puncta per square micrometer. Projected density maps were smoothed for display with a 1 *µ*m Gaussian kernel.

#### Statistical Analysis

Statistical analyses were performed either using MATLAB R2025b or GraphPad Prism (version 10.4.1). Normality was determined by the KS test, and equal variances were checked using Bartlett’s test. If there were more than three groups, the data were normally distributed, and the variances were equal, an ANOVA with multiple comparisons (Tukey’s honestly significant difference procedure). If the data in three or more groups were not normal, the Kruskal-Wallis test with pairwise Mann-Whitney U tests was automatically performed without the need to check for equal variances. In comparisons involving two groups, if the data were normally distributed and the variances were equal, a student’s t-test was conducted. If the data in the two groups were normally distributed, but the variances were unequal, Welch’s t-test was performed. If the data in either group were not normally distributed, a Mann-Whitney U-test was performed to determine statistical significance.

## 1 Data and Code Availability

Raw data is available upon reasonable request. The analysis algorithm for balance beam performance quantification is available in the BEAMCROSS GitHub page.

## Acknowledgements

We are grateful for the help and support provided by the Technology Animal Resources section of Core Facilities at Okinawa Institute of Science and Technology Graduate University for facility maintenance and animal care.

## Author contributions statement

T.T., M.U., and B.I. conceived and designed the experiments, T.T. conducted the behavioral experiments, immunohistochemical stainings, and data acquisition. H.H supported immunohistochemical stainings and animal maintenance. S.D. conducted immunohistochemical stainings for Cx36. T.S. contributed to the conceptualization of the neuropil analysis pipeline. T.T. and M.Y. analyzed the results, prepared the figures, and wrote the initial manuscript. T.T. drew all schematic illustrations using Adobe Illustrator 2026. All authors reviewed the manuscript.

